# All-Aqueous Printing of Viscoelastic Droplets in Yield-Stress Fluids

**DOI:** 10.1101/2022.09.14.507847

**Authors:** Jinchang Zhu, Li-Heng Cai

**Author notes:** Correspondence: L.-H. Cai, **Corresponding author contact:**, Dr. Li-Heng Cai, 228 Wilsdorf Hall, University of Virginia, 395 McCormick Road, Charlottesville, VA 22904.

## Abstract

All-aqueous printing of viscoelastic droplets (aaPVD) in yield-stress fluids is the core of an emerging voxelated bioprinting technology that enables the digital assembly of spherical bio-ink particles (DASP) to create functional tissue mimics. However, the mechanism of aaPVD is largely unknown. Here, by quantifying the dynamics of the whole printing process in real-time, we identify two parameters critical to aaPVD: (1) acceleration of print nozzle, and (2) droplet/nozzle diameter ratio. Moreover, we distinguish three stages associated with aaPVD: droplet generation, detachment, and relaxation. To generate a droplet of good roundness, the ink should be a highly viscous shear-thinning fluid. Using particle image velocimetry and scaling theory, we establish a universal description for the droplet displacements at various printing conditions. Along the direction of nozzle movement, the droplet displacement is determined by the detachment number, a dimensionless parameter defined as the ratio between the dragging force from the nozzle and the confinement force from the supporting matrix. Perpendicular to the direction of nozzle movement, the droplet displacement is determined by the Oldroyd number, a dimensionless parameter that describes the yielded area of the supporting matrix near the print nozzle. For a relaxed droplet, the droplet tail length is independent of droplet/nozzle diameter ratio but determined by the nozzle acceleration. We conclude that printing droplets of good fidelity requires a relatively large droplet/nozzle diameter ratio and intermediate nozzle accelerations. These ensure that the droplet is more solid-like to not flow with the nozzle to form a tadpole-like morphology and that the confinement force from the yield-stress fluid is large enough to prevent large droplet displacement. Our results provide the knowledge and tools for *in situ* generating and depositing highly viscoelastic droplets of good roundness at prescribed locations in 3D space, which help establish the foundational science for voxelated bioprinting.

**Statement of Significance:** Analogues of pixels to two-dimensional (2D) pictures, voxels – in the form of small cubes or spheres – are the basic units of three-dimensional (3D) objects. All-aqueous printing of viscoelastic droplets (aaPVD) is the core of voxelated bioprinting, an emerging technology that uses spherical bio-ink voxels as building blocks to create 3D tissue mimics. By quantifying the dynamics of the whole printing process in real-time, we distinguish the stages associated with aaPVD, identify parameters critical to aaPVD, and develop a universal understanding for *in situ* generating and depositing viscoelastic droplets of good roundness at prescribed locations in 3D space. Our results help establish the foundational science for voxelated bioprinting and provide a general approach for precisely manipulating viscoelastic voxels in 3D space.

## 1. Introduction

Unlike simple Newtonian fluids such as water, bio-inks are complex non-Newtonian fluids that contain biopolymers and cells. Manipulation of such viscoelastic fluids is the core of three-dimensional (3D) bioprinting technology. For instance, in extrusion-based bioprinting, one of the most used 3D bioprinting techniques because of simple operation and bio-friendly processing conditions, bio-ink filaments are extruded from the print nozzle and assembled layer-by-layer to manufacture tissue mimics [1,2]. Yet one-dimensional (1D) filaments are not the basic building blocks of 3D structures. By contrast, analogous of pixels to two-dimensional (2D) pictures, voxels – in the form of either small cubes or spherical particles – are the basic units of 3D objects [3]. In principle, the location, composition, and properties of individual voxels and voxel-voxel interactions can be precisely defined to match the artistry of biological tissues. Such voxelated bioprinting would dramatically expand the capability of existing 3D bioprinting technologies but requires on-demand generation, position, and assembly of each voxel in 3D space.

Precise manipulation of viscoelastic voxels represents both a fundamental and a technological challenge in soft matter science and 3D bioprinting. For example, monodisperse emulsions or droplets can be generated by exploiting Rayleigh–Plateau instability, during which a liquid thread breaks into droplets to minimize their surface area and thus interfacial free energy [4]. This phenomenon forms the scientific foundation of many important technologies such as inkjet-based (bio)printing [5], electrohydrodynamic spraying [6], and droplet-based microfluidics [7–9]. Unfortunately, these technologies have virtually no control over the position of droplets in 3D space. This limitation can be partially circumvented by embedded droplet printing [10–13], which generates and disperses aqueous droplets into an immiscible continuous oil phase made by yield-stress fluids. Because the yield-stress fluid reversibly transitions from solid-like to liquid-like at critical stress, the fluid can stabilize and entrap a droplet once it forms, providing somewhat of control over the droplet position in 3D space. Nevertheless, the breakup of the dispersing fluid thread inevitably forms discrete droplets, which can be used to generate patterns with prescribed droplet distance but precludes controlled assembly of the droplets. Alternatively, using lipid molecules to coat water-in-oil droplets provides an approach to assemble droplets [14–16]. As two lipid-coated droplets come together, the hydrophobic tails of the lipid molecules stack to form a lipid bilayer that cohesively joins the two droplets without coalescence. This process enables creating a droplet network in which aqueous droplets are compartmentalized by the network of lipid bilayers. However, because the lipid bilayer is stabilized by van der Waals force, the network is mechanically weak and cannot support complex structures. Moreover, all these technologies exploit the same mechanism – Rayleigh-Plateau instability – to generate droplets. Consequently, they can handle low viscosity liquids only and must involve organic solvents that are often biohazardous.

Instead of using *in situ* generated viscoelastic bio-ink droplets, using pre-made solid-like particles as building blocks circumvents the technological challenges associated with the on-demand generation of viscoelastic voxels. A prominent example is the printing of tissue spheroids, 3D cell aggregates that better emulate organs compared to traditional 2D cell cultures [17,18]. Although delicate, the spheroids are solid-like and can be successfully manipulated with great care. For instance, a spheroid can be gently picked up by a glass pipette under controlled vacuum aspiration, transferred to a supporting hydrogel matrix, and deposited at a prescribed location upon vacuum removal [19,20]. Through cell migration, closely placed spheroids can fuse to form microtissues with pre-defined shapes. Thus, 3D bioprinting of spheroids resembles some of the features required for voxelated bioprinting but is limited to pre-made solid-like voxels.

Very recently, built on the advancement of embedded 3D printing [21–24], we proved the concept of a voxelated bioprinting technology that enables the digital assembly of spherical particles (DASP) [25]. Unlike existing embedded droplet printing techniques that can only disperse low viscosity fluids in oil-based supporting matrices, DASP generates a highly viscoelastic bio-ink droplet in an aqueous yield-stress fluid, deposits the droplet at a prescribed location, and assembles individual droplets through controlled polymer swelling. Consequently, DASP represents a class of embedded bioprinting technology that enables on-demand generation, position, and assembly of highly viscoelastic voxels in an aqueous environment. Using DASP, we created mechanically robust, multiscale porous scaffolds composed of interconnected yet distinguishable hydrogel particles for responsive insulin release. Despite the practical success, it has yet to be elucidated the mechanism of DASP. *First*, how to generate a droplet with a good roundness without the help from large interfacial tension? *Second*, what are the mechanisms for detaching the print nozzle from a viscoelastic droplet? These questions cannot be answered by applying existing knowledge for printing droplets of low viscosity Newtonian liquids in an immiscible organic fluid.

Here, we systematically explore the mechanism for all-aqueous printing of viscoelastic droplets (aaPVD) in yield-stress fluids. We build a 3D printing platform that allows for exquisite control over the printing conditions and for real-time quantification of the printing dynamics. We identify two parameters critical to aaPVD: (1) acceleration, *a*, of the print nozzle, and (2) the ratio between droplet and nozzle diameters, *D_d_/D_n_*. We distinguish *three* stages associated with aaPVD: (i) extrude the ink to generate a droplet, (ii) detach the print nozzle from the droplet, and (iii) allow the detached droplet to relax (**Fig. 1**). We show that to generate a droplet of good roundness the ink must be a highly viscous, shear-thinning fluid. Using particle image velocimetry (PIV) measurements and scaling arguments, we establish a universal description for the droplet displacement at various combinations of printing conditions. For a relaxed droplet, we discover that droplet tail length is independent of *D_d_/D_n_* but determined by the nozzle acceleration *a* only. We conclude that printing droplets of good fidelity requires that (i) *D_d_/D_n_* should be relatively large and (ii) the nozzle should be detached from the droplet at intermediate accelerations. These ensure that the droplet is more solid-like to not flow with the nozzle to form a tadpole-like morphology, and that the confinement force from the supporting matrix is large enough to prevent large droplet displacement. Our results provide the knowledge and tools for *in situ* generating and depositing highly viscoelastic droplets of good roundness at prescribed locations in 3D space, which help establish the foundational science for voxelated bioprinting.

**Fig 1.**
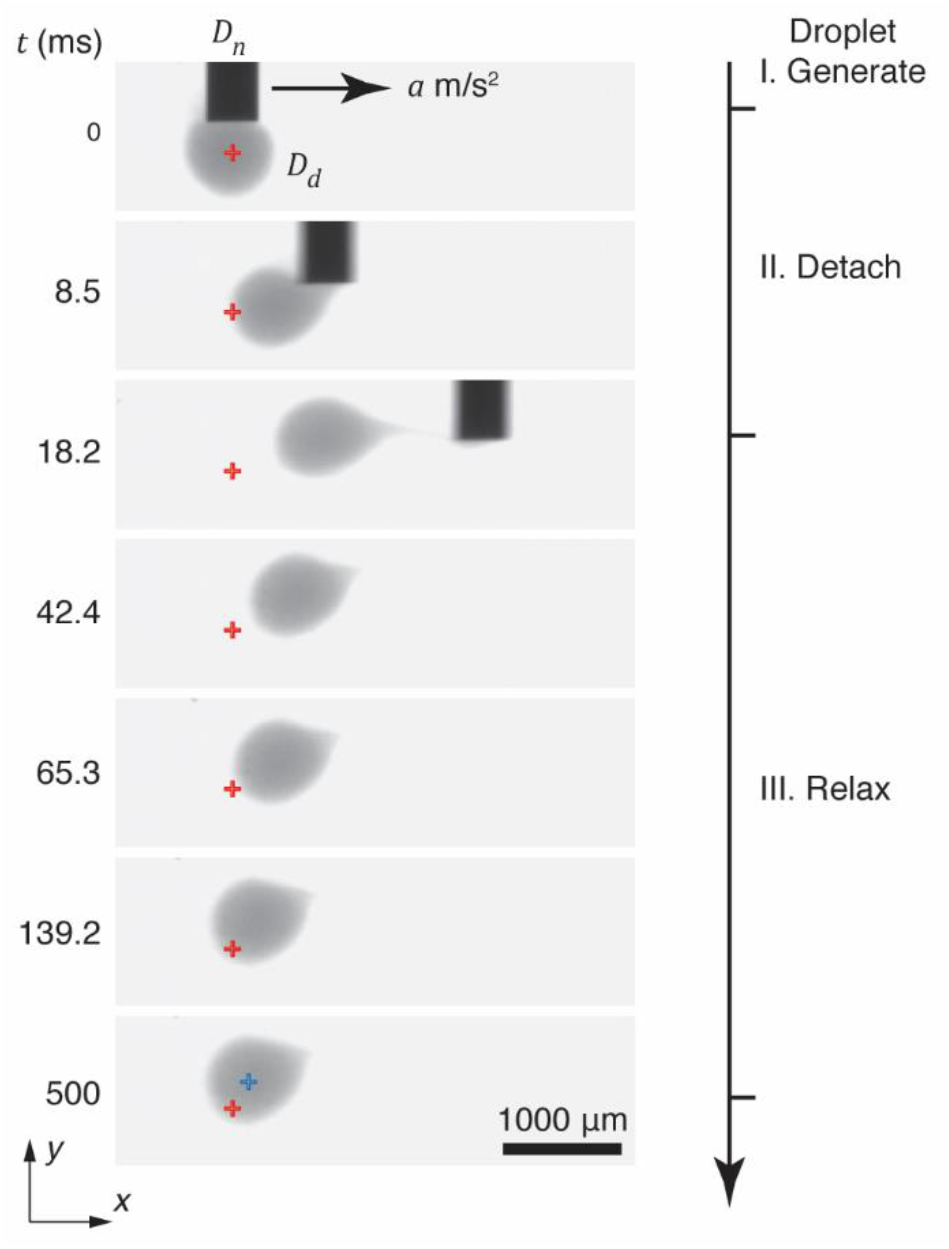
Real-time visualization of printing a viscoelastic droplet in a yield-stress fluid. The printing process consists of three stages: (i) generate a droplet by extruding the bio-ink in the yield-stress fluid, (ii) detach the print nozzle from the droplet, and (iii) allow the droplet to relax from deformation attributed to nozzle movement. The fidelity of droplet printing is defined by two parameters: (1) droplet roundness at the generation stage and after relaxation, and (2) droplet displacement after relaxation. Red cross: center of the droplet at generation stage; blue cross: center of the droplet after relaxation. *D_n_*, nozzle tip diameter; *D_d_*, droplet diameter; *a*, acceleration of the nozzle.

## 2. Materials and Methods

### Materials and reagents

Alginic acid sodium salt (medium viscosity, Cat. No. A2033) and poly(ethylene oxide) (PEO) with MW of 1000 kDa (Cat. No. 372781) are purchased from Sigma-Aldrich (USA). Raw honey is purchased from Private Selection®(USA). Poly(acrylic acid), Carbomer 940 (Cat. No. THK-CARBM-01) is purchased from MakingCosmetics Inc. (USA). 4- (2-hydroxyethyl)-1-piperazineethanesulfonic acid (HEPES, 1M solution, Cat. No. J16924-AP) is purchased from Thermo Scientific (USA). Silica beads (Cat. No. G4649) for particle imaging velocimetry (PIV) are purchased from Sigma-Aldrich (USA). All chemicals are used as received unless otherwise specified.

### Carbomer supporting matrix and inks

To prepare the supporting matrix, we mix 150 mg Carbomer powder with 45 mL DI water and stir the mixture overnight at room temperature to obtain a homogenous solution. Because poly(acrylic acid) carries a large number of carboxylic acid groups, the solution is acidic rather than neutral. Consequently, the rheological properties of Carbomer are sensitive to pH [23]. To this end, we add 5 mL 4-(2-hydroxyethyl)-1-piperazineethanesulfonic acid (HEPES), a non-ionic amphoteric buffer, to precisely control the pH of the supporting matrix at 6.75. During this process, the supporting matrix changes from an opaque to an optically transparent hydrogel with the final concentration of Carbomer 0.3% (w/v) and HEPES 100 mM. The hydrogel is centrifuged at 3000 rpm to remove bubbles before use. To prepare the alginate-based inks, we dissolve alginic acid sodium salt in DI water at 4.0% (w/v), and then sonicate the mixture for 2 h at 60 *°C* to make a homogenous solution. The alginate solution is stained by food color with a volume ratio of 1% to contrast the supporting matrix. Raw honey is used without any treatment for the extrusion study. The hybrid bio-ink, 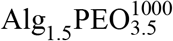, is prepared by dissolving alginate and PEO with MW of 1000 kDa sequentially in DI water 1.5% and 3.5% (w/v), at respectively. All bio-inks are stored in a 4 °C refrigerator before use.

### Instrumentation of droplet printing

We build the platform for droplet printing by integrating an extrusion module into a linear driving stage customized based on Aerotech AGS1500 gantry. The system allows positioning of the print nozzle with an accuracy of ±1.5 μm on both X and Y axes and 10 μm on Z axis. Moreover, the system can drive the print nozzle up to the speed of 3 m/s at the acceleration up to 2.5 g, allowing us to explore a wide range of printing conditions. The extrusion module is built based on a linear screw (T8) actuator which converts the rotary motion of a stepper motor (Aerotech, SM35) into translational motion. We use the extrusion module to drive a glass syringe (1 mL, Shanghai Bolige Industrial & Trade Co., Ltd) with a speed and acceleration specified by the controlling software of the printing system. To visualize the process of droplet printing in real-time, we integrate a long working distance objective into a high-speed camera (FASTEC, IL5) that is connected to and controlled by a computer.

### Rheometry

Rheological measurements are performed using a stress-controlled rheometer (Anton Paar, MCR 302) equipped with parallel-plate geometries of diameter 50 mm with rough surface to avoid the slippage between hydrogels and the geometry. To measure the shear moduli of the supporting matrix and inks, we conduct a frequency sweep from 0.1 to 100 Hz at an oscillatory shear strain of 0.5%. To characterize the yield-stress behavior of the supporting matrix, we conduct a stress sweep from 0.1 to 1000 Pa at an oscillatory frequency of 1 Hz. To precisely determine the yield stress, we use a previously developed method [26] by fixing the stress while monitoring the shear rate until reaching a steady state. By progressively increasing the stress level, we obtain the dependence of stress on the steady shear rate. To characterize the self-healing behavior of the supporting matrix, we apply a periodic destructive high shear strain and monitor the mechanical properties of the supporting matrix in real-time. For each cycle, we instantly increase the shear strain to 1000% within 1 sec, and then apply an oscillatory shear strain of 1% at a frequency of 1 Hz for 200 seconds, during which both *G’* and *G”* are measured. To characterize the dependence of viscosity of inks on shear rate *γ*, we conduct a shear rate sweep from 10^-3^ to 100 1/sec, during which the shear stress and viscosity are recorded.

### Droplet printing

We print a droplet in the supporting matrix contained in a glass container with the dimension of 75×25×25 mm. We use the same print nozzle (26G, McMaster-Carr) with an inner diameter *D_n,i_* = 300 μm and an outer diameter of *D_n_* = 440 μm for all experiments unless otherwise specified. The tip of the nozzle is placed 10 mm below the surface of the supporting matrix and at least 10 mm away from the wall of the container; these distances are more than 10 times of the droplet diameter, such that the boundary effects on the droplet printing can be ignored. In all printing experiments, we extrude the ink at a fixed rate to create a droplet of prescribed volume, and then detach the print nozzle from the droplet at a prescribed acceleration. During droplet detachment, we move the print nozzle by at least 40 mm, which is no less than 40 times of the droplet diameter, to ensure that the print nozzle is completely detached from the droplet. Simultaneously, we use a fast camera operated at 500 frames per second to monitor the whole printing process including droplet extrusion, detachment, and relaxation.

### Droplet tracking

We use FIJI TrackMate to obtain the trajectory of a droplet during the whole printing process. The droplet is recognized by Laplacian of Gaussian (LoG) filter, which is a derivative filter that detects the rapid intensity change at the edge of the droplet. However, because the intensity also changes rapidly near the edge of the nozzle, some spots can be mistakenly recognized by the LoG filter; this would cause errors in correlating identified spots in neighboring frames to generate the droplet trajectory. To avoid this, we implement Linear Assignment Problem (LAP) tracker to enumerate all possible correlations and select the one with the highest possibility to exclude the mistakenly recognized spots. Finally, we overlay the generated droplet trajectory with the corresponding movie to validate the accuracy of droplet tracking.

### Particle image velocimetry (PIV) measurements

To visualize the motion of supporting matrix, we disperse silica beads with an average diameter of 50 μm in the supporting matrix. The Stokes number, defined as *St* = *t_b_/t_n_*, where *t_b_* is the bead response time and *t_n_* is the time scale of nozzle movement, is on the order of 10^-11^, as 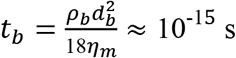 and 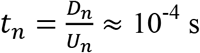, where *ρ_b_* = 2.65 g/cm^3^ is the density of silica beads, *d_b_* = 50 μm is the bead diameter, *η_m_* is the dynamic viscosity of the matrix with the value on the order of 10 Pa·s (**Fig. S2**), *D_n_* = 440 μm is the nozzle outer diameter, and *U_n_* ≈ 2 m/s is the maximum speed of the nozzle. Therefore, the beads can be assumed to follow the motion of the supporting matrix nearly perfectly. In addition, we control the concentration of the beads at 4 mg/mL, corresponding to a volume fraction of 1.5×10^-3^; this small value ensures that the effects of particles as fillers on the flow properties of the supporting matrix is negligible. Moreover, the silica beads have a refractive index 1.48 that is higher than that of the supporting matrix 1.33; this contrast allows the visualization of the beads using a fast camera under bright field, as shown by the small white dots in **Movie S2** and **Movie S11**. For every image, the scaling factor is 8.46 μm/pixel; therefore, each bead is about 6 pixels, a number recommended by PIV analysis. Moreover, the sharp contrast between the tracer beads and the matrix allows us to use contrast-limited adaptive histogram equalization (CLAHE) filter to obtain a binary image for subsequent PIV analysis.

To obtain the velocity vectors, we implement 2D cross-correlations of the bead distribution within a chosen interrogation window [27]. We start with the coarse interrogation windows of 128×128 pixels with 50% overlap to estimate and offset the noise from lens shake, and then set the fine interrogation windows of 64×64 pixels with 50% overlap to calculate the velocity field. The size of the interrogation window has no direct influence on the spatial resolution. Yet, for particle images larger than the 3 pixels, the random error of PIV measurements is typically 0.05 pixels for a given interrogation domain [28]. Moreover, because the displacement reflects an average displacement of all particle images in the interrogation window, it requires an optimized number of particles per sample area. In our PIV measurements, we use about 6 particles per interrogation window, a number recommended to ensure reliable PIV analysis. Furthermore, as recommended by PIV analysis, the frame steps are chosen to ensure that the displacement peak in the cross-correlation function is within the range of 10 – 20 pixels [29]. All image processing and calculations are performed using MATLAB (PIVlab) [30] unless otherwise specified.

### Statistical analysis

All particle trajectories are shown as mean with sample size *n*=3. All data points are shown as mean ± S.D. with sample size *n*=3.

## 3. Results and Discussion

To explore the mechanisms of droplet printing, we develop a 3D printing platform consisting of *four* modules (**Fig. S1**). *First*, unlike typical desktop 3D printers lacking precise control over the motion profile of the print nozzle, we customize a high-precision linear-driving stage that enables precise control over the moving speed and acceleration of the print nozzle. Specifically, the nozzle can move a resolution of 1.5 μm at accelerations over nearly two orders of magnitude from 0.1 to 25 m/s^2^; this allows for probing time-scale dependent viscoelastic properties of inks and supporting matrix. *Second*, to extrude a viscoelastic droplet with a prescribed volume at precisely controlled speed, we integrate into the positioning system a linear driving stage to mechanically extrude viscoelastic bio-inks. *Third*, we use an optically transparent yield-stress fluid made of 0.3% (w/v) poly(acrylic acid) hydrogel (Carbomer 940) as the supporting matrix; this allows for direct visualization of the whole printing process [23]. Because poly(acrylic acid) is a charged polymer, we use HEPES buffer to control the pH at 6.75 to ensure stable rheological properties (see **Materials Methods**). Using oscillatory shear measurements, we confirm that the supporting matrix transitions from an elastic solid with a shear modulus of 75 Pa to a liquid above a critical shear stress (**Fig. 2A&B**). Quantitatively, the yield-stress behavior can be described by Herschel-Bulkley model [31]:

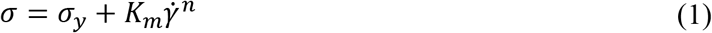

in which *σ* is the shear stress, 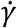 is shear rate, *σ_y_* = 9.5 Pa is the yield stress, *K_m_* = 4.0 Pa·s^n^ is the consistency index of the matrix, and *n* = 0.45 in the flow index (**Fig. 2C**). Moreover, the supporting matrix can self-heal rapidly within 1 sec for unlimited times (**Fig. 2D**). These features are critical to supporting matrix for embedded 3D printing, as established in our previous studies [25]. *Fourth*, we customize a long-working distance objective that can be mounted to a fast camera to visualize the whole printing process in real-time (see **Materials and Methods**). Together, our 3D printing platform allows for exquisite control over printing conditions and real-time quantification of the printing dynamics, providing the tools necessary for elucidating the mechanisms of all-aqueous droplet printing.

**Fig 2.**
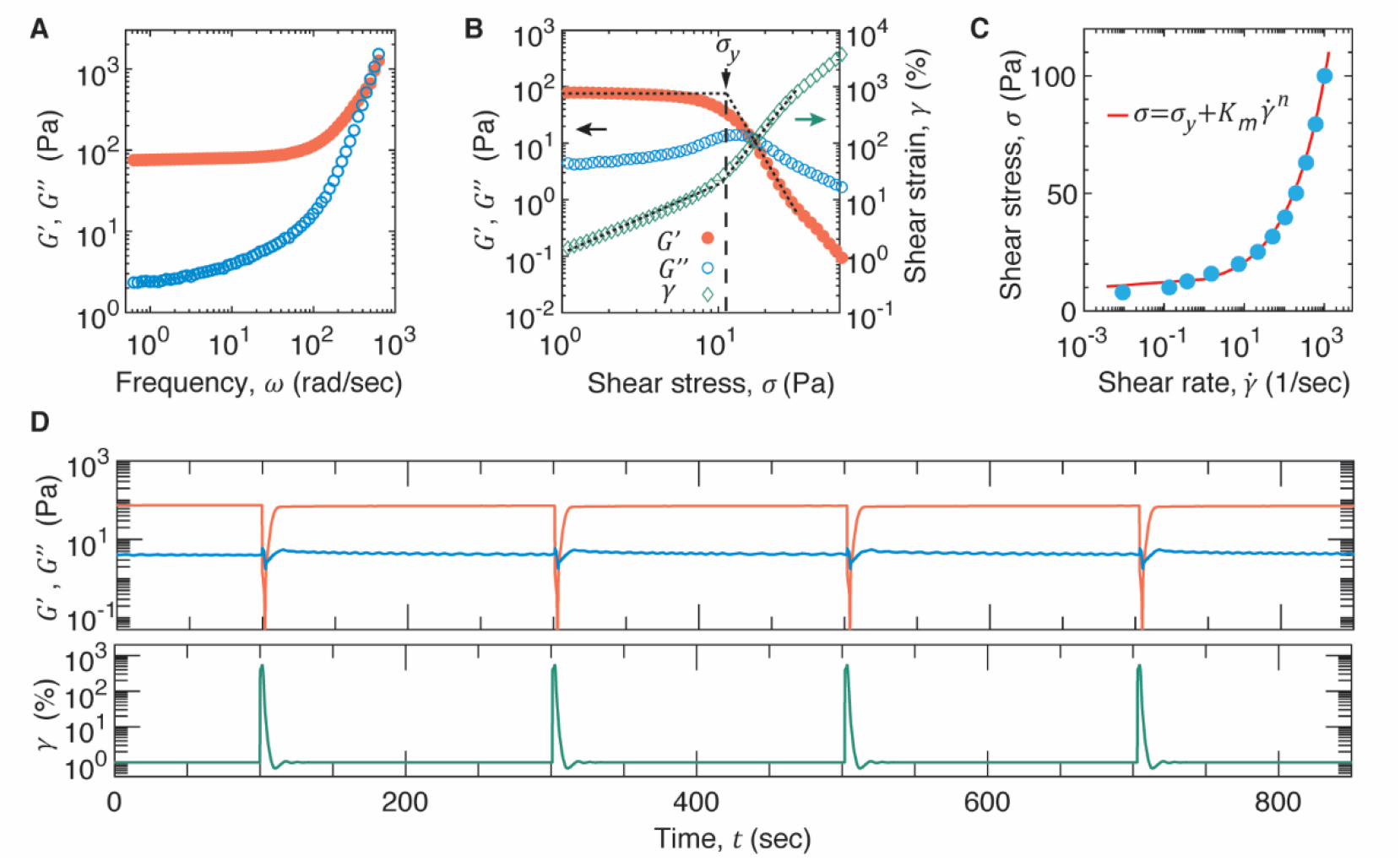
Physical properties of supporting matrix. The supporting matrix is made of 0.3% (w/v) Carbomer in HEPES buffer. (**A**) Dependence of the storage (filled circles, *G’*) and loss (empty circles, *G”*) moduli of the supporting matrix on oscillatory shear frequency *ω* measured at a fixed strain of 0.5%at 20 °C. (**B**) The Carbomer supporting matrix is a yield-stress fluid that becomes fluid-like above the yield stress *σ_y_*. Left *y*-axis: the dependence of *G’* and *G”* on shear stress *σ*. Right *y*-axis: the dependence of shear strain (empty squares), *γ*, on shear stress. The measurements are performed at oscillatory frequency of 1 Hz at 20 °C. (**C**) Dependence of the shear stress for the supporting matrix on shear rate 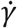. Circles are experimental data measured by controlling the shear stress while measuring the steady shear rate. Solid line is the fit to the experimental data using Herschel-Bulkley model 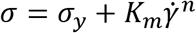 [see eq. (1)], in which *σ_y_* = 9.5 Pa is the yield stress, *K_m_* = 4.0 Pa·s^n^ is the consistency index of the matrix, and *n =* 0.45 in the flow index. (**D**) The supporting matrix self-heals to recover its original properties for unlimited times. Upper: Real-time behavior of *G*’(red) and *G”*(blue) under a periodic destructive high shear strain of 1000%. The shear strain is applied for 1 s, and the waiting time between two neighboring destructive shear strains is 200 s.

We divide the whole printing process into *three* stages based on the moving profile of the print nozzle. In a typical printing process, we *first* position the print nozzle at a prescribed location to extrude a droplet with a prescribed diameter *D_d_*, *then* detach the nozzle from the droplet at a controlled acceleration *a*, and *finally* wait for the droplet to relax, as shown by the image series in **Fig. 1**. To describe the fidelity of droplet printing, we introduce two parameters: (1) droplet roundness at the generation stage and after relaxation, and (2) final droplet displacement after relaxation (red and blue crosses in **Fig. 1**).

### 3.1 Droplet generation

To explore the mechanism of droplet generation, we choose three fluids as inks with representative flow properties: (1) water – a Newtonian fluid of low viscosity 1 mPa·s; (2) honey – a Newtonian fluid of high viscosity 25 Pa·s that is nearly constant over a wide range of shear rate from 10^-3^ to 10^2^ s^-1^ (red triangles in **Fig. 3A**) and (3) an alginate solution – a non-Newtonian, shear thinning fluid whose viscosity decreases with the increase of shear rate (blue circles in **Fig. 3A**). We tune the concentration of alginate to be 4.0% w/v (Alg_4.0_), such that its low shear rate viscosity is about 40 Pa·s, close to that of honey (**Fig. 3A**). Yet, at shear rates higher than 0.5 s^-1^, the apparent viscosity of the alginate ink decreases with the shear rate by a power law:

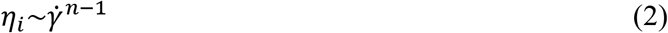

in which with the flow index *n* = 0.26 ± 0.08, as shown by the dashed line at the right of **Fig. 3A**.

**Fig 3.**
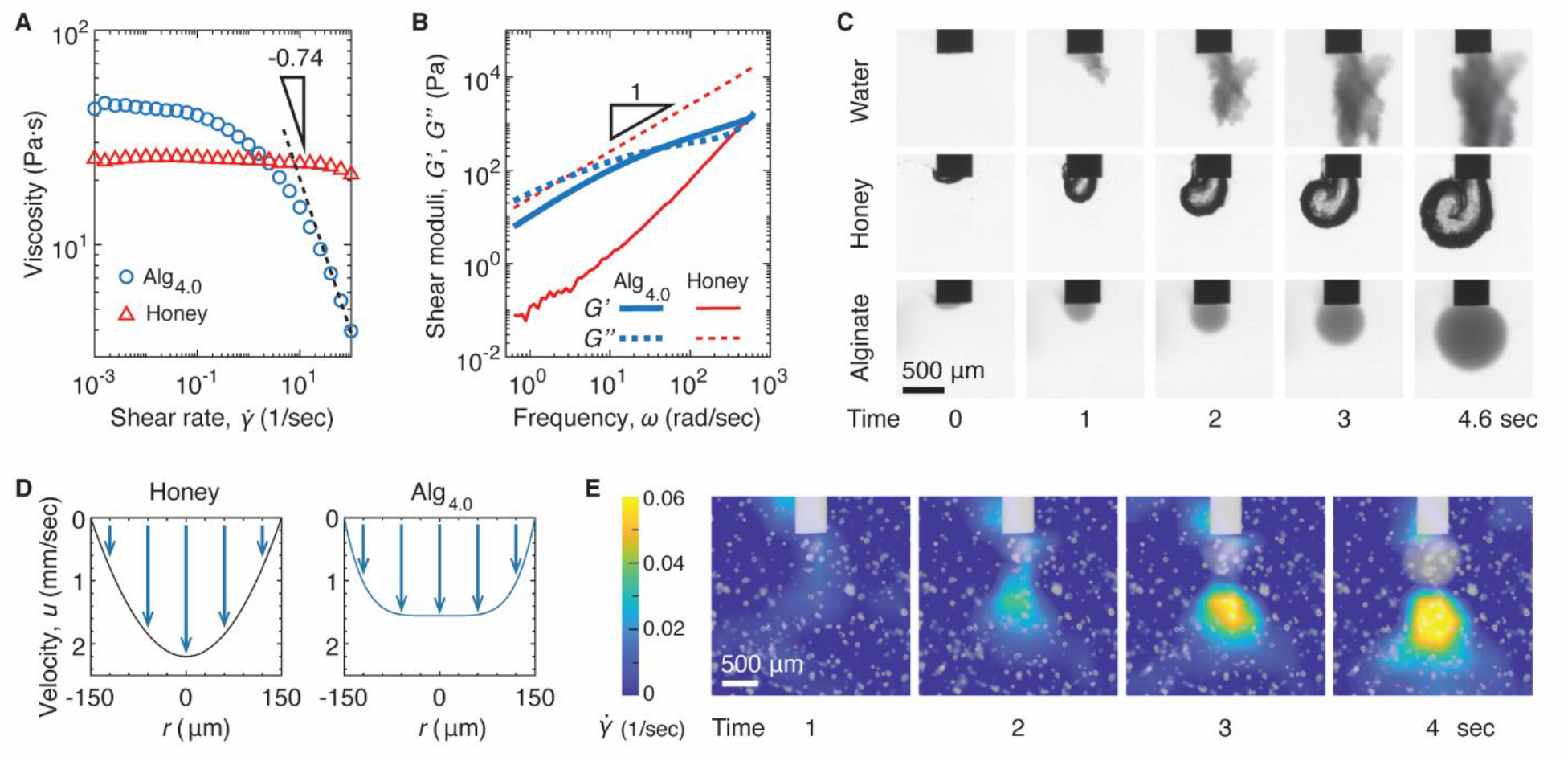
Extruding droplets with good roundness requires viscous, shear-thinning inks. (**A**) Dependence of viscosity of inks on shear rate 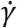. At high shear rates, the alginate ink is a power-law fluid [see eq. (2)]. Alg_4.0_ denotes an alginate solution with the concentration of 4% w/v. (**B**) Dependence of shear storage (solid line, *G’*) and loss (dash line, *G”*) moduli on oscillatory shear frequency at a fixed strain of 0.5%at 20 °C. (**C**) Real-time visualization for extruding droplets of inks of low viscosity water, high viscosity honey, and viscoelastic alginate solution (4.0% w/v) in a yield-stress fluid made of carbomer (**Fig. 2**). (**D**) Velocity profile of the viscous, Newtonian like honey and the shear-thinning alginate ink flowing through the cylindrical nozzle. (**E**) An example PIV analysis that shows the strain rate field of the supporting matrix during the extrusion stage (**Movie S2**). Nozzle outer diameter, *D_n_* = 440 μm; droplet diameter, *D_d_* = 880 μm. White dots: 50 μm silica beads that are used as tracers for PIV analysis.

The flow properties of the inks are further confirmed by oscillatory shear frequency measurements. For instance, the loss modulus *G”* of honey increases linearly with frequency, confirming that honey is a Newtonian fluid (red dashed line in **Fig. 3B**). By contrast, for Alg_4.0_ the storage modulus *G’* becomes higher than loss modulus *G”* at high shear frequency, a phenomenon characteristic of a viscoelastic liquid. Importantly, all these three inks are soluble in water. These features make the three liquids ideal candidates for exploring the rheological properties of inks required for generating droplets of good roundness in yield-stress fluids.

We use the same print nozzle to extrude the inks at a constant flow rate to generate droplets. We fix the inner diameter of the nozzle *D_n,i_* at 300 μm and extrude an ink at a constant volumetric flow rate of *Q* = 0.08 μL/sec, which corresponds to an average flow velocity *U* = 1.1×10^-3^ m/sec and a shear rate of 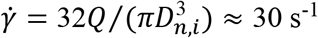. The ink is mixed with food dyes (see **Materials and Methods**) to generate contrast in color with the supporting matrix, such that we can use a fast camera to monitor the process of droplet extrusion in real time.

When extruding the low viscosity water, the flow flushes out of the nozzle and forms an irregular, branched pattern, as visualized by the optical images in the top panel of **Fig. 3C**. This behavior is in sharp contrast to extruding water into oil-based supporting matrix, where a thread of water forms droplets to minimize interfacial energy [10–13]. The observed branched pattern is reminiscent of viscous fingering, or Saffman-Taylor instability [32]: when a low viscosity fluid pushes a more viscous fluid, the interface between the two fluids develops an instability that leads to the formation of fingerlike patterns [26]. Yet, different from Saffman-Taylor instability where the finger-like patterns develop along the directions of the flow, the front of the water flow ramifies randomly (top right panel in **Fig. 3C**). This is because, compared to the yield stress *σ_y_* = 9.5 Pa of the supporting matrix (**Fig. 2C**), the shear stress associated with extruding the low viscosity water is more than two orders of magnitude smaller, 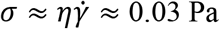, in which *η* = 10^-3^ Pa·s is the water viscosity. Therefore, the water flow cannot mechanically displace the supporting matrix. Moreover, because of negligible interfacial tension between water and the supporting matrix, the water easily flows through the micrometer scale interparticle space of the porous supporting matrix, as evidenced by the water flow in **Movie S1**. These results show that low viscosity liquids are not suitable for generating droplets in porous yield-stress fluids without the help of large interfacial tension.

Extruding a way much more viscous honey does not lead to random ramification but a droplet-like pattern (middle panel, **Fig. 3C**). Similarly, extruding highly viscous Alg_4.0_ forms a droplet (bottom panel, **Fig. 3C**). These are because the shear stress associated with extruding these highly viscous inks, 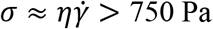, is much higher than the yield stress 9.5 Pa of the supporting matrix. Therefore, instead of penetrating through the porous supporting matrix, the flow of highly viscous fluid mechanically displaces the supporting matrix. Indeed, PIV measurement reveals a large strain field of the supporting matrix beneath the droplet (**Fig. 3E**).

However, unlike honey which does not develop to a round droplet but forms a curly strip (middle panel, **Fig. 3C**), Alg_4.0_ ink forms a nearly perfect spherical droplet. This is likely attributed to the difference in the velocity profile of fluids flowing through the cylindrical nozzle. The velocity of a flow stream decreases with the increase of the distance *r* from the center of the nozzle [33]:

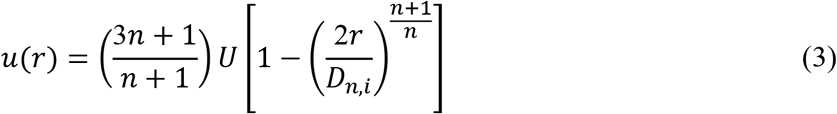

in which *U* is the average velocity, and *n* if the flow index. For Newtonian fluids like honey, the flow index *n =* 1; whereas for the shear thinning fluid Alg_4.0_, the flow index is very small, *n* = 0.26 [see eq. (2)]. Thus, compared to the Newtonian-like honey, the velocity profile of the shear-thinning Alg_4.0_ is much flatter, as illustrated in **Fig. 3D**. This results in a more uniform displacement of the supporting matrix, as shown by the relatively flat strain field at the forefront of the droplet in **Fig. 3E** and **Movie S2**. Consequently, the droplet grows more uniformly, as shown in real-time by **Movie S1**. Consistent with this understanding, extruding another highly viscous shear-thinning ink results in droplets of good roundness (**Fig. S3**). Collectively, our results indicate that a highly viscous, shear-thinning ink is suitable for generating a droplet of good roundness in yield-stress fluids.

### 3.2 Droplet detachment

Using droplets as building blocks to create complex structures requires each droplet to be printed at a prescribed location. This requires detaching the print nozzle without significantly displacing the droplet. To understand the mechanism of droplet detachment, we explore the dependence of droplet displacement on nozzle acceleration, one of the most important parameters of droplet printing. We fix the nozzle diameter at *D_n_* = 440 μm, extrude an alginate droplet (Alg_4.0_) with a fixed diameter of *D_d_* = 880 μm, and detach the print nozzle from the droplet at various accelerations. Simultaneously, we use a fast camera to monitor the droplet motion. Because of the geometric symmetry of the cylindrical nozzle and the spherical droplet, we focus on the side view of the printing process, as exemplified by the time-series photographs in **Fig. 4A** and **Movie S7**.

**Fig 4.**
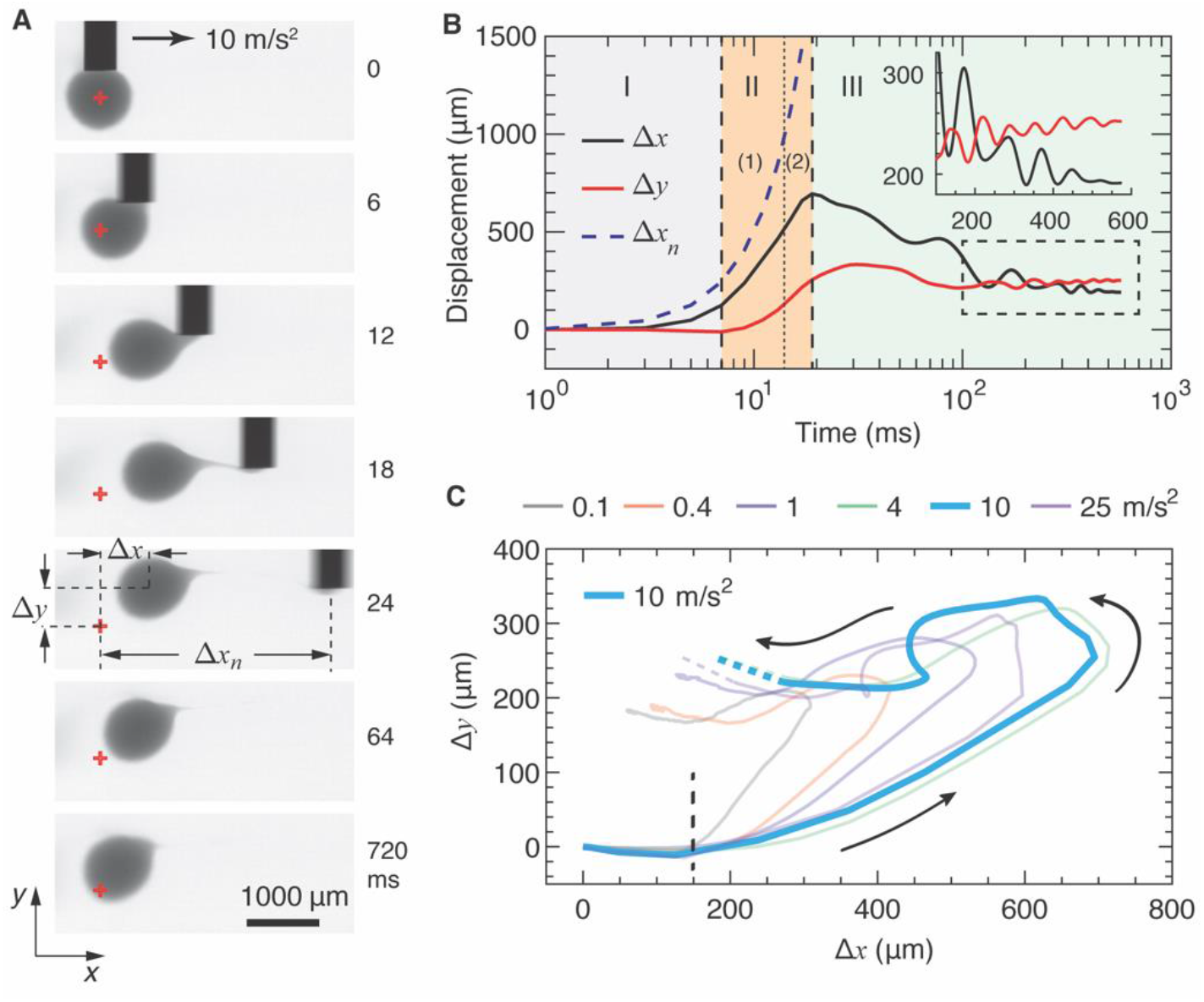
The trajectory of a droplet during the detachment stage. (**A**) A representative time-series of photographs for detaching the print nozzle from a droplet at acceleration *a* = 10 m/s^2^ (**Movie S7**). Red cross: center of the droplet at generation stage. (**B**) Dependence of the displacements of the print nozzle and the droplet on time at the printing conditions in (**A**). Δ*x_n_* is the displacement of the nozzle; Δ*x* and Δ*y* are, respectively, the displacements of the droplet along the moving direction of the nozzle and along the nozzle axis. Insert: Droplet displacement after 100 sec. (**C**) Trajectories of droplets printed at various accelerations. Dashed lines: extrapolated trajectories based on final position of the droplet. In all experiments, the nozzle diameter is fixed at *D_n_* = 440 μm, and the droplet diameter is fixed at *D_d_* = 880 μm. Trajectories are shown as mean with sample size *n*=3

#### 3.2.1 Droplet trajectory

Using the movie for *a* = 10 m/s^2^ as an example, we distinguish *three* regimes based on the moving profile of the droplet and its position relative to the nozzle. Initially, the droplet moves along the direction of nozzle movement (*x* axis) by a distance Δ*x*_1_ (*Regime I*, **Fig. 4B**). Afterward, the droplet moves not only along *x* axis but also upward along the length of nozzle (*y* axis) (*Regime II*, **Fig. 4B**). Yet initially the droplet remains to be attached to the nozzle (*Regime II-1*); then, the droplet is separated from the nozzle but continues the moving trend (*Regime II-2*). During this process, a thin, long string of the viscoelastic droplet material is pulled out along the moving direction of the nozzle, as visualized by the photograph at 18ms in **Fig. 4A**. At the end of *Regime II*, the droplet reaches its maximum displacement, Δ*x_m_* and Δ*y_m_*. Finally, the droplet moves backward but does not completely return to its initial position with a non-zero final displacement of Δ*x_f_* and Δ*y_f_* (*Regime III*, **Fig. 4B**)). During this process, the string of viscoelastic droplet material partially springs back. These three regimes are found for a wide range of accelerations ranging from 0.1 m/s^2^ to 25 m/s^2^, as shown by the trajectories in **Fig. 4C** and **Movies S3**-**S8**.

To better understand the process of droplet detachment, we quantify the droplet displacement at various accelerations in each regime. In *Regime I* the droplet displacement Δ*x*_1_ is nearly a constant 180 *μ* m regardless of accelerations, as indicated by the vertical dashed line in **Fig. 4C**. Simultaneously, the print nozzle moves by nearly the same distance, suggesting that the droplet moves along with the nozzle. Yet, because the droplet is not a solid particle but liquid-like, there exists a small lag between droplet and nozzle displacements. In *Regime II*, the droplet reaches its maximum displacement, which exhibits a nonmonotonic dependence on the acceleration. As *a* increases, Δ*x_m_* increases by more than twice from 308 μm at 0.1 m/s^2^ to 694 μm at 10 m/s^2^ and then decreases to 596 μm at 25 m/s^2^. Similar trend is observed for Δ*y_m_*. *In Regime III*, as the droplet starts to return to its initial position, we identify two types of motion. At relatively *low* accelerations below 1 m/s^2^, the droplet moves backward following a smooth trajectory. By contrast, at *high* accelerations above 4 m/s^2^, the droplet moves backward following an oscillatory trajectory (**Fig. 4C**). This is likely because that under high accelerations the print nozzle inputs a large amount of energy into the supporting matrix, such that the matrix must oscillate to dissipate the energy. Indeed, the embedded droplet oscillates along both *x* and *y* directions with a damped magnitude but nearly the same periods, as shown by the black and red curves in *Regime III* and by the inset in **Fig. 4B**. As the supporting matrix relaxes, the droplet reaches its final position. Yet, the droplet does not return to its initial position regardless of accelerations, as shown by the end points of trajectories in **Fig. 4C**. These results suggest that detaching the print nozzle inevitably displaces the droplet to some extent.

#### 3.2.2 Final droplet displacement

As a droplet moves through the supporting matrix, the droplet experiences friction that increases with the droplet diameter. Therefore, in addition to nozzle acceleration, it is expected that the droplet size influences the final displacement of a droplet. To explore this, we quantify the final displacements, Δ*x_f_* and Δ*y_f_*, for droplets of various sizes (**Movies S7, S9, S10**). We fix the nozzle outer diameter at *D_n_* = 440 μm and change the droplet diameter *D_d_* from 440, 660 to 880 μm, which correspond to the droplet/nozzle diameter ratios of *D_d_/D_n_* =1.0, 1.5, and 2.0, respectively. As expected, the final displacement increases with the decrease of droplet size, as visualized in **Fig. 4A** and **Fig. 5A**. Similar trends persist for various nozzle accelerations from 0.1 to 25 m/s^2^ (**Fig. 5B**). Remarkably, regardless of the acceleration, the relative increase in the final droplet displacement is nearly the same: as *D_d_/D_n_* decreases from 2 to 1.0, Δ*X_f_* increases dramatically by about 10 times (**Fig. 5B, i**), whereas Δ*y_f_* increases by about 2 times (**Fig. 5B, ii**).

**Fig 5.**
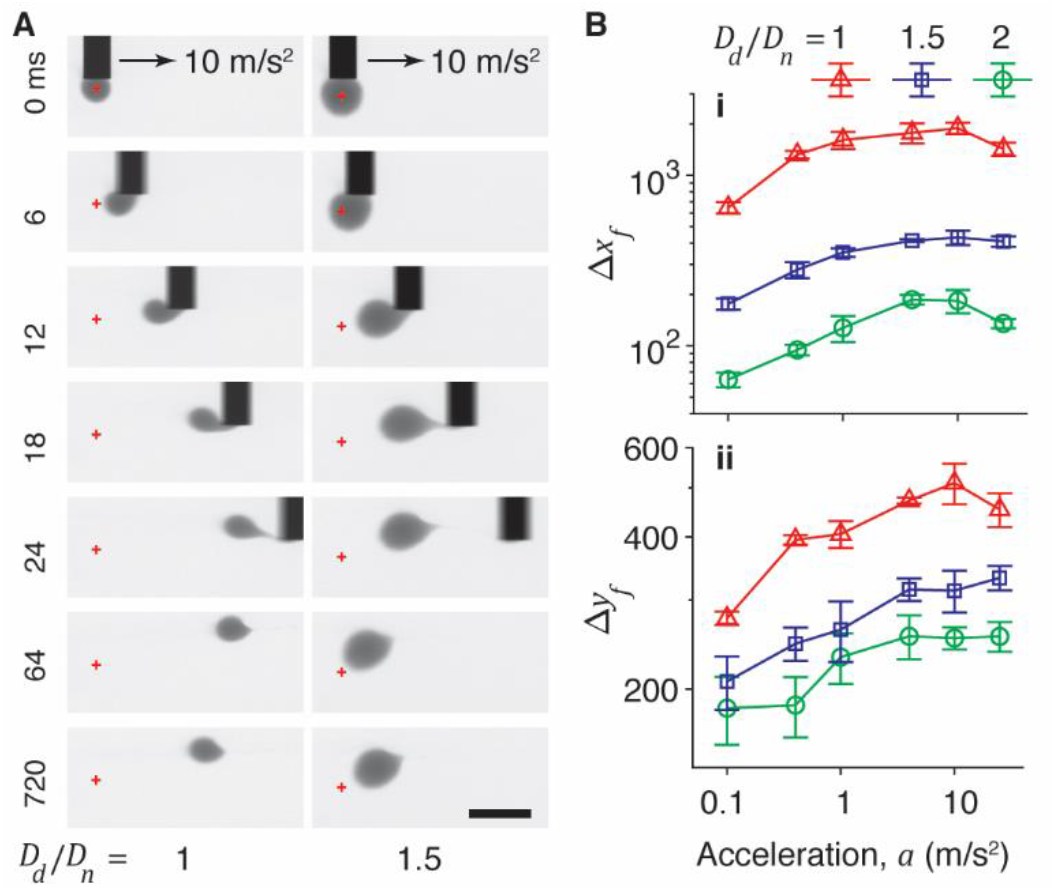
Final displacement of a droplet is determined by nozzle acceleration and droplet size. (**A**) A time-series of photographs for detaching a nozzle from a droplet at a fixed acceleration rate *a* = 10 m/s^2^ but different droplet/nozzle diameter ratio, *D_d_/D_n_*. The nozzle diameter is fixed at *D_n_* = 440 μm, yet the droplet diameter *D_d_* varies from 440 μm (left column, **Movie S9**) to 660 μm (right column, **Movie S10**). Red cross: the center of the droplet at generation stage. Scale bar, 1000 μm. (**B**) The final displacement of droplets at various combinations of *D_d_/D_n_* and nozzle acceleration *a*: (i) the displacement along the moving direction of nozzle, Δ*Xf*; (ii) the displacement along the nozzle axis, Δ*y_f_*. Error bar: STD, *n*=3.

Interestingly, the final droplet displacement exhibits a nonmonotonic dependence on the nozzle acceleration. For instance, at *D_d_/D_n_* = 2.0, the Δ*x_f_* increases from 55 μm at 0.1 m/s^2^ to 186 μm at 4 m/s^2^ and then decreases to 135 μm at 25 m/s^2^ (**Fig. 4C** and green circles in **Fig. 5B,i**). By contrast, Δ*y_f_* increases monotonically by nearly 50% from 190 μm from 260 μm as *a* increases from 0.1 to 4 m/s^2^ but saturates at higher accelerations (green circles in **Fig. 5B, ii**). Moreover, similar trends are observed for different *D_d_/D_n_* values: both Δ*x_f_* and Δ*y_f_* initially increase with *a* but either saturate or slightly decrease at high *a* (**Fig. 5B**). These results contrast the droplet displacement in a viscous fluid, which would increase if the droplet were pulled at higher accelerations or by larger forces. Nevertheless, our results show that the final displacements of the droplet are highly correlated to both the nozzle acceleration and droplet diameter.

To further understand the origin of final droplet displacements, we perform PIV measurements to quantify the deformation of supporting matrix during the detachment process (see **Materials and Methods**). We discover that the flow of the supporting matrix is highly correlated to the trajectory of the droplet. Initially, as the nozzle accelerates, the matrix mainly flows along the direction of nozzle movement (*x* direction) (<10 ms in **Movie S11**); this corresponds to the droplet trajectory in *Regime I* where the droplet displaces along *x* direction only (**Fig. 4B**). Then, the matrix continues to flow along with the nozzle, but starts to flow upward along *y* direction to fill the empty space left behind the nozzle, as visualized by the photograph at 10 ms in **Fig. 6A**. Simultaneously, the droplet rotates by nearly 90°, as shown by the photographs of droplets in **Fig. 6B**. This behavior is consistent with the droplet trajectory in *Regime II* where the droplet moves along both *x* and *y* directions (**Fig. 4B**). Finally, as the nozzle is separated from the droplet and continuing to move away from the droplet, the matrix bounces back through a damping process, during which the matrix largely oscillates along *x* direction to recover from the deformation caused by the nozzle movement, as shown by the photographs >20 ms in **Fig. 6A** and by **Movie S11**. The observed velocity fields of the matrix are highly consistent with the damping trajectory of the droplet in *Regime III* (**Fig. 4B)**.

**Fig 6.**
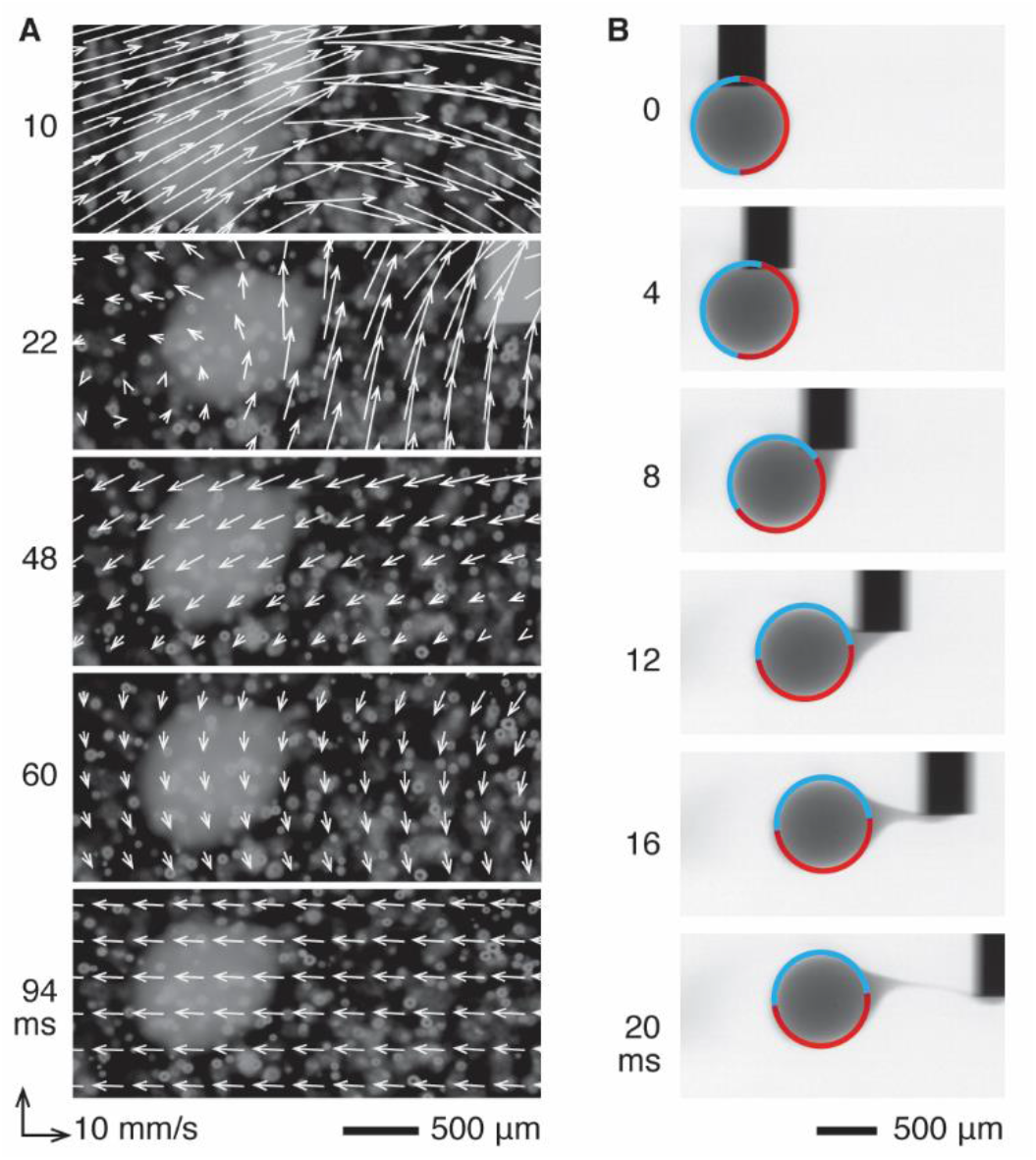
Velocity field of the supporting matrix, droplet rotation, and droplet displacement during the detachment stage. (**A**) An example PIV analysis that shows the velocity field of the supporting matrix during the droplet detachment stage (**Movie S11**). Nozzle tip diameter, *D_n_* = 440 μm; droplet diameter, *D_d_* = 880 μm. White dots: 50 μm silica beads that are used as tracers for PIV analysis. White arrows denote the absolute velocity field of the supporting matrix near the droplet. (**B**) A time-series of photographs show that a droplet rotates and displaces under the dragging from the nozzle.

The correlation between matrix flow and droplet trajectory indicates that the droplet displacement consists of recoverable and irrecoverable parts. The recoverable displacement is determined by the reversible, elastic deformation of the supporting matrix, during which the droplet moves together with the surrounding matrix flow. This recoverable displacement is associated with *Regimes II-2* and *III* when the nozzle has been completely detached from the droplet (**Fig. 4B**). By contrast, the irrecoverable displacement is determined by the irreversible, plastic deformation of the supporting matrix, during which the droplet not only is dragged by the nozzle but also moves along with the plastic flow of the supporting matrix near the nozzle. Importantly, the plastic flow is attributed to the rearrangement of microgel particles, which occurs after microgel particles being yielded and displaced by the nozzle. Such an irrecoverable droplet displacement is associated with *Regimes I* and *II-1* where the nozzle remains in contact with the droplet (**Fig. 4B**).

#### 3.2.3 Scaling analysis for final droplet displacement

Based on the above understanding, we develop a scaling theory to describe the dependence of final droplet displacement on printing parameters. For the final displacement along the movement of nozzle, Δ*x_f_*, we identify two forces exerted to the droplet during the detachment process: dragging force *f_d_* from the nozzle, and confinement force *f_c_* from the supporting matrix, as illustrated in **Fig. 7A**. The dragging force is determined by fracturing the connection between the nozzle tip and the droplet, which is proportional to the product of the contact area, 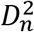, and the effective shear modulus of alginate ink, *G_i,e_*:

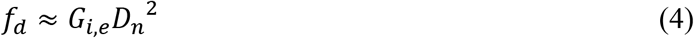

**Fig 7.**
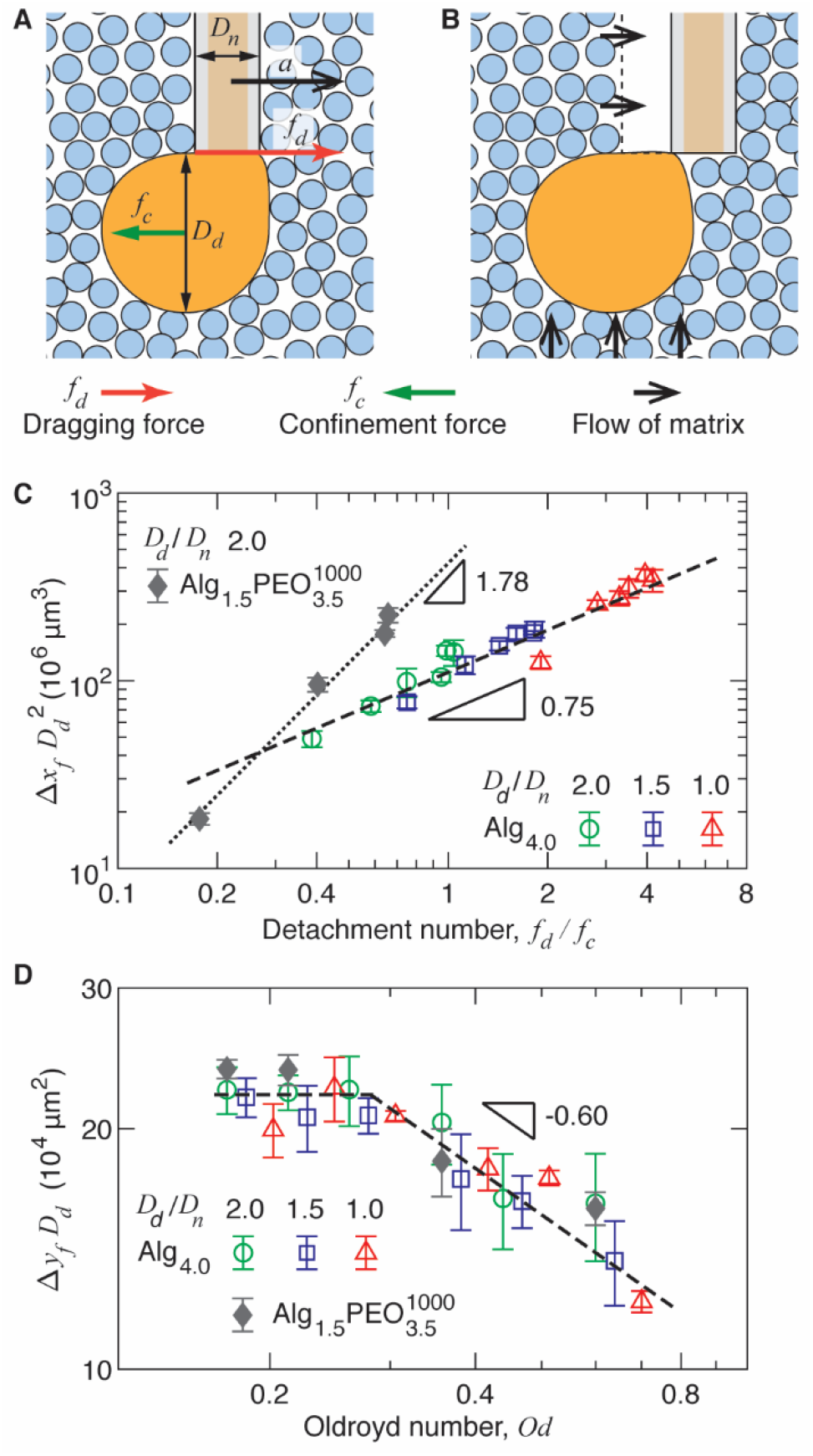
Universal behavior of droplet final displacement. (**A**) As the print nozzle detaches from the droplet, the droplet experiences two competing forces: (1) the dragging force associated with separating the print nozzle from the droplet, and (2) confinement force from the supporting matrix exerted on the droplet. (**B**) The moving nozzle leaves an “empty” space behind, which will be filled by the supporting matrix nearby. The flow of the supporting matrix pushes the droplet along *y* axis. (**C**) The displacement volume of the droplet along the movement of nozzle, Δ*x_f_D_d_*^2^, scales with the detachment number that is defined as the ratio between the dragging and the confinement forces, 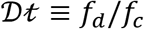 [see eq. (8)], by a power of 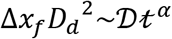. For ink Alg_4.0_, *α* = 0.75 (empty symbols); for ink Alg_1.5_PEO_3.5_^1000^, *α* = 1.78 (filled diamonds).. (**D**) The dependence of cross-area traveled by the droplet along *y* axis, Δ*y_f_D_d_*, on the Oldroyd number [see eq. (10)]. Error bar: STD, *n*=3.

Unlike a particle moving in viscous liquids that experiences friction determined by the fluid viscosity, in our experiments the droplet is confined in a yield-stress fluid, which is effectively an elastic solid below the yield stress. Therefore, the droplet is expected to be mechanically constrained in the supporting matrix. The confinement force *f_c_* is the product of the droplet cross area, *D_d_*^2^, and the effective modulus of the matrix, *G_m,e_*:

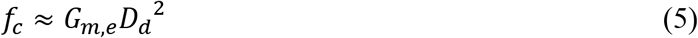

We emphasize that the *G_m,e_* is not the equilibrium shear modulus of the supporting matrix; instead, it is the modulus of the supporting matrix experienced by the droplet if it were moving along with the nozzle.

Because both the supporting matrix and the alginate ink are viscoelastic fluids, their effective moduli are dependent on the probing time scale, *τ_p_*. To estimate *τ_p_*, we consider the moving profile of the print nozzle during the detachment stage. Unlike existing embedded droplet printing techniques in which the nozzle is moving at a constant speed [12], in our experiments the nozzle moves at a constant acceleration *a*. Therefore, the probing time scale associated with our droplet printing is:

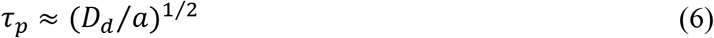

Alternatively, the probing time scale can be understood in the context of materials deformation rate. As the print nozzle accelerates, it deforms the droplet at a rate of 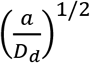. In the meantime, the droplet tends to displace the supporting matrix at an acceleration of *a*, reminiscent of inertia force from Newton’s second law. Consequently, the supporting matrix is effectively deformed by the droplet at a rate of 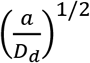, which is the reciprocal of the probing time scale [see eq. (6)].

We determine the effective moduli at the probing time scale based on the experimentally measured shear storage moduli from oscillatory shear measurements (**Fig. 2A** and **Fig. 3A**). The probing time scale *τ_p_* is the inverse of the angular frequency or rotation rate of the geometry, *ω* ≈ 1/*τ_p_*; therefore, we can directly map the shear storage moduli for both the ink and the supporting matrix at each printing condition. This allows us to determine the ratio between the dragging force and the confinement force applied to the droplet, which is defined as the *detachment number, Dt*:

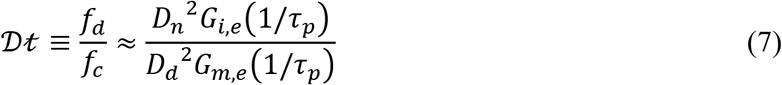

Because *f_d_/f_c_* describes the amount of energy input by the print nozzle, we plot the final *displacement volume* of the droplet along the nozzle movement, Δ*x_f_D_d_*^2^, against *f_d_/f_c_*; this collapses all the data points regardless of printing conditions, as shown by the symbols in **Fig. 7C**. Moreover, this universal dependence can be described by a simple power law relation:

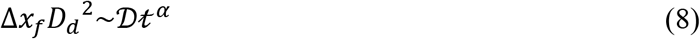

in which the scaling exponent *α* = 0.75±0.03 for Alg_4.0_, as shown by the dashed line in **Fig. 7C**. This universal scaling relation suggests that Δ*x_f_* can be reduced by decreasing the detachment number 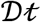.

Unlike Δ*x_f_* that is highly dependent on the dragging force, along *y* axis the droplet does not experience the dragging force from the print nozzle. However, the competition between the dragging force and the confinement force generates a torque, as illustrated by the pair of green and red arrows in **Fig. 7A**. This torque results in the rotation of the droplet, as experimentally verified in **Fig. 6B**. Importantly, as the nozzle translates, it leaves a conceptually empty space behind the nozzle, as schematically illustrated in **Fig. 7B**. This space must be filled by the supporting matrix near the print nozzle, as illustrated by black arrows in **Fig. 7B**. Thus, the droplet will move along with the matrix flow; this effectively results in the displacement of the droplet along *y* direction. Indeed, this phenomenon is experimentally verified by PIV analysis, as shown by the velocity fields of the supporting matrix and the droplet position at 10 ms in **Fig. 6A** and by the PIV analysis of the whole process in **Movie S11**. These results suggest that Δ*y_f_* is highly correlated to flow of the supporting matrix, which is largely determined yielded area near the print nozzle.

The *yielded area* near the nozzle can be characterized by the Oldroyd number 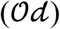 [34], which is defined as the ratio of the yield stress of the supporting matrix to the viscous stress generated by the print nozzle: 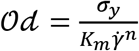, in which *σ_y_* = 9.5 Pa is the yield stress, *K_m_* = 4.0 Pa·s^n^ is the consistency index of the matrix, and *n* = 0.45 in the flow index [see eq. (1)]. The shear rate of the supporting matrix equals the ratio between the nozzle velocity *U_n_* and the nozzle diameter: 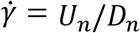. Because the nozzle moves at constant acceleration *a*, we use the average velocity when the nozzle moves by the droplet diameter *D_d_*, beyond which the nozzle starts to separate from the droplet: *U_n_* ≈ (*D_d_a*)^1/2^. Thus, for a given set of printing conditions (nozzle diameter, droplet diameter, and nozzle acceleration), the Oldroyd number is:

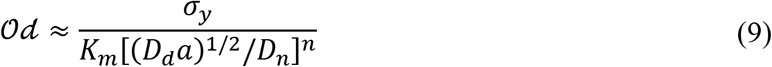

Because the Oldroyd number characterizes the yielded area near the print nozzle, we plot the *cross area* traveled by the droplet along *y* axis, Δ*y_f_D_d_*, against the Oldroyd number. Remarkably, this collapses all the data sets to a master curve, as shown by the symbols in **Fig. 7D**. Moreover, we identify two regimes for the dependence of Δ*y_f_D_d_* on the Oldroyd number:

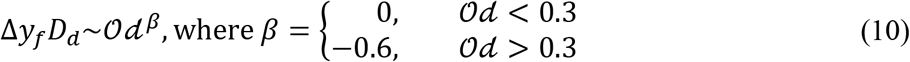

For small values with 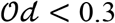, Δ*y_f_D_d_* remains nearly constant. By contrast, at relatively large values with 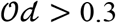, Δ*y_f_D_d_* decreases with the increase of Oldroyd number by a power of −0.6 (dashed lines in **Fig. 7D**). The universal scaling relation between the droplet displacement area and the Oldroyd number suggests that Δ*y_f_* can be reduced by decreasing the Oldroyd number.

The scaling relations predict that, for the same supporting matrix, the detachment number depends on the rheological properties of the ink [eq. (8)], whereas the Oldroyd number is independent of the ink [eq. (10)]. To test these predictions, we exploit the concept of hybrid bio-inks [25] to develop an additional ink, Alg_1.5_PEO_3.5_^1000^, which is a polymer solution consisting of 1.5% (w/v) alginate and 3.5% (w/v) 1000 kg/mol poly(ethylene oxide) (PEO). This ink has a low-shear rate viscosity comparable to that of Alg_4.0_ but is more elastic with a smaller loss factor, tan *δ* = *G”/G’*, because of the long PEO polymers (**Fig. S3A-C**). Using this ink, we print droplets at various nozzle accelerations; example trajectories are shown in **Fig. S3D** and time-lapsed photos for the printing process are shown in **Fig. S3E**. Remarkably, similar scaling relations are observed. Yet, the displacement volume along the *x-*axis scales with the detachment number by a power of 1.79 (filled diamonds in **Fig. 7B**), much larger than 0.75 for Alg_4.0_. This behavior is consistent with the understanding that at the same printing conditions more elastic droplets tend to be dragged by a larger distance. By contrast, the displacement along the *y*-axis follows a master curve the same as that for Alg_4.0_ (filled diamonds in **Fig. 7C**); this behavior further supports the understanding that Δ*y_f_* is determined by the matrix but not so much by the ink. Taken together, our results indicate that the developed scaling relations provide a universal description for droplet displacements. Importantly, the scaling relations guide how to experimentally control the final droplet displacements along with both *x* and *y* directions. The methods for reducing both Δ*x_f_* and Δ*y_f_* are the same: (1) increasing droplet to nozzle diameter ratio *D_d_/D_n_*, (2) increasing the yield-stress and the stiffness of the supporting matrix, and/or (3) using relatively low nozzle acceleration.

### 3.3 Droplet relaxation

Embedded droplet printing requires depositing a droplet not only at prescribed location but also with a good roundness. The morphology of a relaxed droplet (*Stage III* in **Fig. 1A**) is largely determined by the detachment stage, where an extruded droplet is deformed by the print nozzle. During this process, the nozzle shears the droplet and sometimes pulls a string from the droplet to form a tadpole-like morphology, as shown by **Movie S3-S8** and visualized by the photographs in **Fig. 8A**. Moreover, the length of the tail of the tadpole-like pattern decreases with the increase of the nozzle acceleration (**Fig. 8A**). Qualitatively, this behavior is consistent with the understanding that at higher accelerations, or shorter probing time scales, the alginate ink has less time to relax, such that the droplet is more solid-like and less prone to flow with the nozzle.

**Fig 8.**
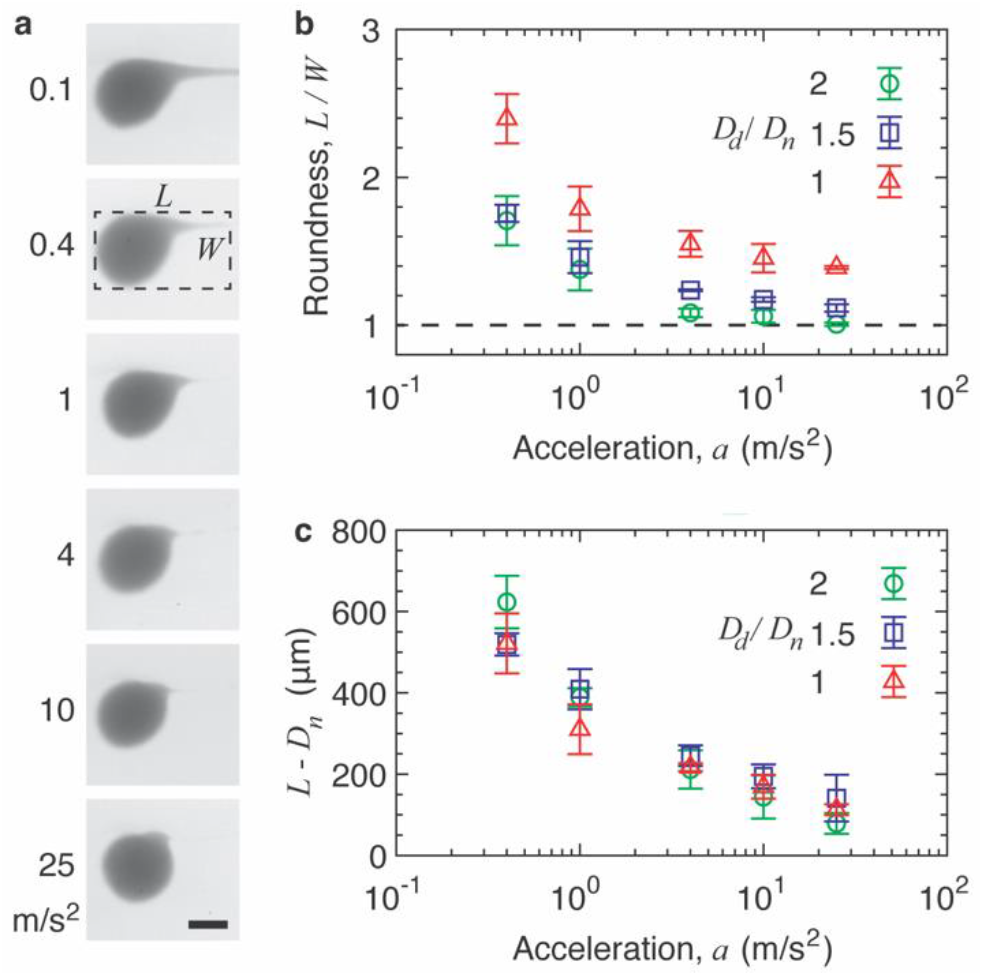
Roundness of a relaxed droplet is determined by the acceleration of print nozzle. (**A**) Representative photographs of relaxed droplets printed at various nozzle accelerations. Nozzle tip diameter, *D_n_* = 440 μm; droplet diameter, *D_d_* = 880 μm. The roundness of a droplet is defined as the ratio between the droplet length along the nozzle movement direction, *L*, and the droplet width along the nozzle axis, 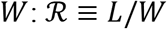. Scale bar, 500 μm. (**B**) Dependence of the droplet roundness on the nozzle acceleration *a*. (**C**) Dependence of the droplet tail length *L* – *D_d_* on the nozzle acceleration. Nozzle tip diameter is fixed at *D_n_* = 440 μm. Error bar: STD, *n*=3.

To describe the morphology of relaxed droplet, we introduce a parameter, droplet roundness 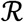, which is defined as the ratio between the length and head width of the tadpole, 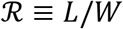, as illustrated in **Fig. 8A.** The larger the value of 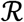, the less round the droplet is. We quantify the roundness of droplets printed at various combinations of nozzle accelerations and droplet sizes. We find that the droplet roundness decreases with increase of the droplet/nozzle diameter ratio *D_d_/D_n_*, as shown by the lower shift from red triangles to green circles in **Fig. 8B**. This suggests that increasing droplet size promotes the formation of a round droplet, enhancing the fidelity of droplet printing.

However, the absolute length of the droplet tail, *L* – *D_d_*, is independent of droplet size but determined by the nozzle acceleration only, as shown by the collapsed data points in **Fig. 8C**. At low nozzle acceleration *a =* 0.1 m/s^2^, the tail is extremely long and is beyond the measurement range (**Movie S3**). As *a* increases by nearly 50 times from 0.4 to25 m/s^2^, the tail length decreases from ~60% to ~5% of the droplet diameter. We note that the droplet tail length becomes longer for a more elastic ink consisting of long polymers (**Fig. S3E**); this suggests that the tail length is also dependent of the rheological properties of the ink. Nevertheless, for Alg_4.0_ there appears to be a crossover acceleration, *a* ≈ 2 m/s^2^, above which the tail length is relatively small and decreases slowly with the increase of nozzle acceleration. At this crossover acceleration, ink is pulled by the print nozzle at a probing time scale, *τ_p_* ≈ (*D_n_/a*)^1/2^ ≈ 10 ms, which is comparable to the relaxation time of the alginate ink, *τ_relax_* ≈ 1/*ω_c_* ≈ 25 ms, where *ω_c_* is the crossover frequency below which *G”* is higher than *G”* (**Fig. 3B**). These results suggest that to print a droplet of good roundness, it requires the nozzle to move at relatively high accelerations to ensure that the probing time scale is shorter than the relaxation time of viscoelastic ink, such the droplet does not have time to relax and flow with the nozzle.

Collectively, our studies show that printing droplets of good fidelity requires: (1) relatively large droplet/nozzle diameter ratio, (2) relatively stiff supporting matrix with high yield stress, and (3) intermediate nozzle acceleration, at which the associated probing time scale is shorter than the relaxation time of the viscoelastic ink but not too short to result in large dragging force.

## 4. Conclusion

In summary, we have systematically investigated mechanisms for all-aqueous printing of viscoelastic droplets in yield-stress fluids. We build a 3D printing platform that allows for exquisite control over the printing conditions and real-time quantification of the printing dynamics. We identify two parameters critical to droplet printing: (1) acceleration *a* of the print nozzle, and (2) the ratio between droplet and nozzle diameters *D_d_/D_n_*. Further, we distinguish *three* stages associated with droplet printing: (i) extrude the ink to generate a droplet, (ii) detach the print nozzle from the droplet, and (iii) allow the detached droplet to relax.

Remarkably, extruding Newtonian fluids of either low or high viscosity does not warrant a droplet of good roundness. Instead, to generate a droplet of good roundness, the ink should be a highly viscous, shear-thinning fluid. The high viscosity ensures that the stress associated with extruding the ink is much larger than the yield stress of the supporting matrix, such that the growth of the droplet can displace the supporting matrix. The shear-thinning behavior ensures a flat velocity profile as the ink flows out of the nozzle, such that the droplet grows uniformly to achieve a good roundness.

By tracking the trajectory of a droplet, we identify *three* regimes for droplet detachment. Initially, the droplet moves along the direction of nozzle movement (*Regime I*); then, the droplet moves along both the direction of the nozzle movement and the nozzle axis (*Regime II*); finally, the droplet moves back towards its initial position (*Regime III*). By quantifying the velocity field of the supporting matrix, we discover that *Regimes I* and *II* are associated with the plastic, irreversible deformation of the yield-stress fluid, whereas *Regime III* is associated with the elastic, reversible deformation of the yield-stress fluid. Based on this knowledge, we develop a scaling theory to describe the dependence of final droplet displacement on printing conditions. Along the moving direction of the nozzle, the droplet displacement volume exhibits a power law relation with the detachment number, which is a dimensionless parameter defined as the ratio between the dragging force from the nozzle and the confinement force from the supporting matrix. Perpendicular to the moving direction of the nozzle, the droplet displacement area is determined by Oldroyd number, a dimensionless parameter that characterizes the yielded area near the print nozzle. For a relaxed droplet, we discover that droplet tail length is independent of *D_d_/D_n_* but determined by the nozzle acceleration *a:* the higher the acceleration, the shorter the tail is.

Although the scaling relations provide a general understanding of the droplet fidelity at various printing conditions [eqs. (8) and (10)], the value of the exponent in the scaling relations depends on the rheological properties of the inks. A mechanistic understanding of the value of the exponent is beyond the scope of this work and will be the subject of future exploration. Nevertheless, our results suggest three major criteria for all-aqueous printing viscoelastic droplets of good roundness at a prescribed location in yield-stress fluids. *First*, a highly viscous, shear-thinning ink is suitable for generating a droplet of good roundness. *Second*, the nozzle must be detached from the droplet at intermediate accelerations such that the associated probing time scale is shorter than the relaxation time of the viscoelastic droplet. This ensures that the droplet is more solid-like to not flow with the nozzle to form a tadpole-like morphology and that the supporting matrix is effectively stiff enough to confine the droplet to prevent large displacement. Because such a criterion is based on the intrinsic relaxation time scales of the droplets, it should not depend on the direction of nozzle movement. For instance, detaching the print nozzle vertically at a relatively small acceleration of 1 m/s^2^ results in a droplet of large final displacement and elongated shape with tadpole-like morphology (**Fig. S4**, **Movie S13**). *Third*, using a relatively large droplet/nozzle diameter ratio further improves the fidelity of droplet printing. This can be achieved using a tapered glass microcapillary as the print nozzle, whose tip diameter can be easily reduced to tens of micrometers [35]. Indeed, using a microcapillary as the print nozzle to increase droplet/nozzle diameter ratio to six enables negligible final droplet displacement (**Fig. S5**).

Our findings have important implications in biomanufacturing and soft matter science. In the context of technology development, unlike existing 3D bioprinting techniques that are largely reply on using 1D filaments as building blocks, aaPVD provides a general approach for *in situ* generating and depositing highly viscoelastic droplets of good roundness at prescribed locations in 3D space. This approach can be readily extended to assemble bio-ink voxels to create highly heterogenous yet tightly organized tissue constructs [36]. In the context of soft matter science, aaPVD does not rely on the classic Rayleigh-Plateau instability to generate droplets; instead, it exploits previously unexplored nonlinear fluid mechanics of complex fluids to manipulate viscoelastic droplets. The developed knowledge and tools not only help establish the foundational science for voxelated bioprinting but also will stimulate new research directions in soft matter and complex fluids.

## Acknowledgement

We thank Yuehui Wang for assembling the 3D motion system.

## Supplementary materials

Supporting Information is available from Online Library or from the author.

