## supporting information for "All-Aqueous Printing of Viscoelastic Droplets in Yield-Stress Fluids"

**This PDF file includes**

- Figure S1. Setup of 3D printing platform.
- Figure S2. Dependence of viscosity of carbomer supporting matrix on shear rate $\dot{\gamma}$.
- Figure S3. Printing viscoelastic droplets of a hybrid ink consisting of alginate and long linear poly(ethylene oxide) (PEO).
- Figure S4. Using tapered capillary nozzle decreases the final displacement of droplet.
- Movie S1. Extrusion of water, honey, and alginate in a yield-stress fluid.
- Movie S2. Strain rate field during the droplet extrusion stage using PIV analysis.
- Movie S3. Detachment of a cylinder nozzle from a droplet at acceleration $a$ =0.1 m/s$^2$ for nozzle tip diameter $D_n$ = 440 µm and droplet diameter $D_d$ = 880 µm.
- Movie S4. $a$ =0.4 m/s$^2$, $D_n$ = 440 µm, $D_d$ = 880 µm.
- Movie S5. $a$ =1 m/s$^2$, $D_n$ = 440 µm, $D_d$ = 880 µm.
- Movie S6. $a$ =4 m/s$^2$, $D_n$ = 440 µm, $D_d$ = 880 µm.
- Movie S7. $a$ =10 m/s$^2$, $D_n$ = 440 µm, $D_d$ = 880 µm.
- Movie S8. $a$ =25 m/s$^2$, $D_n$ = 440 µm, $D_d$ = 880 µm.
- Movie S9. $a$ =10 m/s$^2$, $D_n$ = 440 µm, $D_d$ = 440 µm.
- Movie S10. $a$ =10 m/s$^2$, $D_n$ = 440 µm, $D_d$ = 660 µm.
- Movie S11. Velocity field of the supporting matrix during droplet detachment stage using PIV analysis.
- Movie S12. Detachment of a tapered capillary nozzle from a droplet at acceleration $a$ =10 m/s$^2$ for nozzle tip diameter $D_n$ = 120 µm and droplet diameter $D_d$ = 720 µm.
- Movie S13. Vertical detachment of a cylinder nozzle from a droplet at acceleration $a$ =1 m/s$^2$ for nozzle tip diameter $D_n$ = 440 µm and droplet diameter $D_d$ = 880 µm.

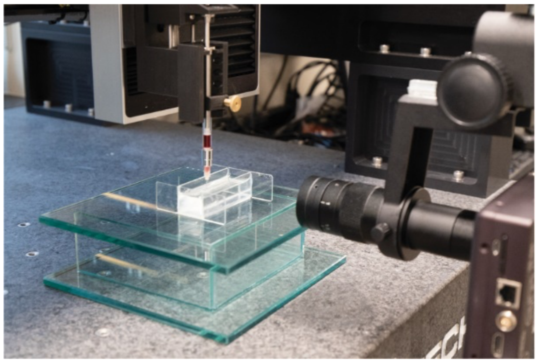

**Figure S1. Setup of 3D printing platform.** A photograph of the 3D motion system mounted with a mechanical extrusion module. The printing process is monitored by a high-speed camera mounted with a long-working distance objective.

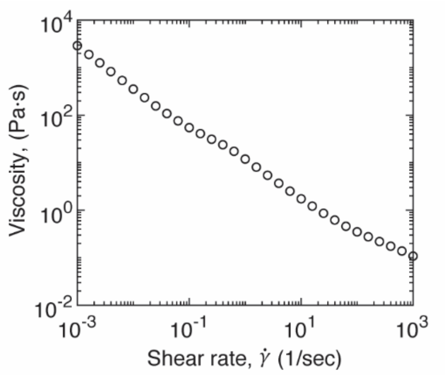


**Figure S2. Dependence of viscosity of carbomer supporting matrix on shear rate $\dot{\gamma}$.**

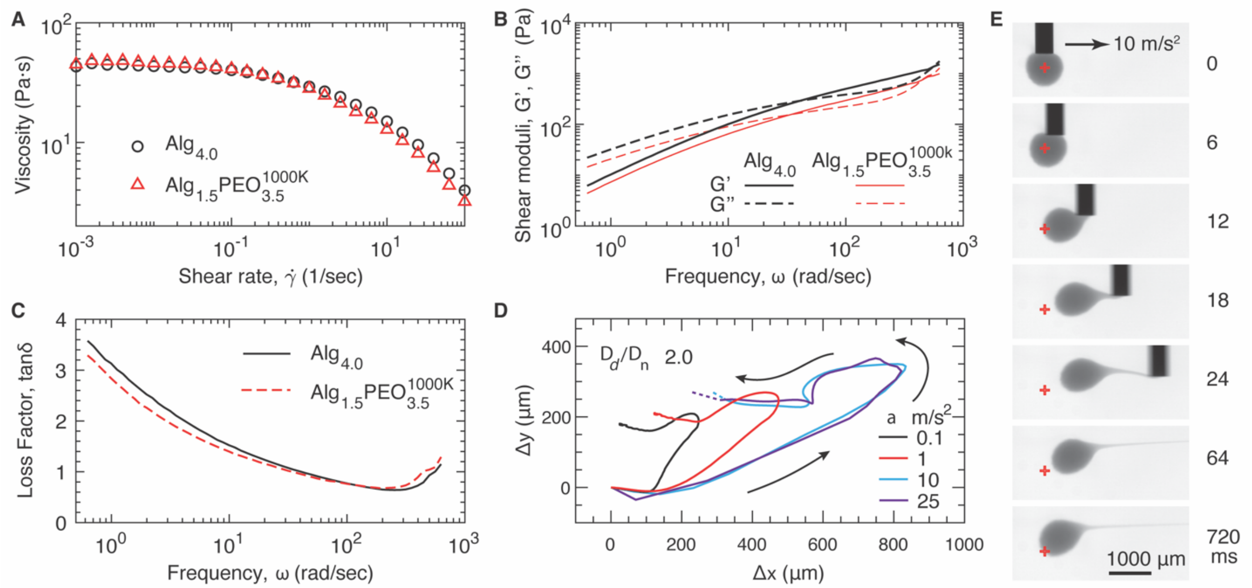


**Figure S3. Printing viscoelastic droplets of a hybrid ink consisting of alginate and long linear poly(ethylene oxide) (PEO).** The ink denoted as $Alg_{1.5}PEO_{3.5}^{1000}$, in which 1.5 and 3.5 are, respectively, the concentration (w/v) of alginate (Alg) and PEO in percentage; 1000, molecular weight PEO in kg/mol. **(A)** Dependence of viscosity on shear rate for $Alg_{4.0}$ and $Alg_{1.5}PEO_{3.5}^{1000}$. **(B)** Dependence of storage ($G'$) and loss ($G''$) moduli of two inks, $Alg_{4.0}$ and $Alg_{1.5}PEO_{3.5}^{1000}$, on oscillatory shear frequency $\omega$ at 0.5% strain and 20 ℃. **(C)** The loss factor, $\tan\delta = G''/G'$, for $Alg_{4.0}$ (black line) and $Alg_{1.5}PEO_{3.5}^{1000}$ (red, dashed line) **(D)** Trajectories of printing droplets of $Alg_{1.5}PEO_{3.5}^{1000}$ ink at various nozzle accelerations $a$. Nozzle diameter, $D_n = 440$ μm; droplet diameter $D_d = 880$ μm. **(E)** A representative time-series of photographs for detaching the print nozzle from a droplet at acceleration $a = 10$ m/s$^2$

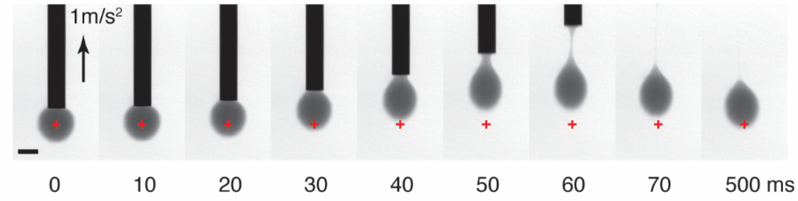


**Figure S4. A representative time-series of photographs for detaching the print nozzle vertically from a droplet.** Acceleration $a = 1$ m/s$^2$. Nozzle diameter, $D_n = 440$ µm; droplet diameter $D_d = 880$ µm. Scale bar, 500 µm.

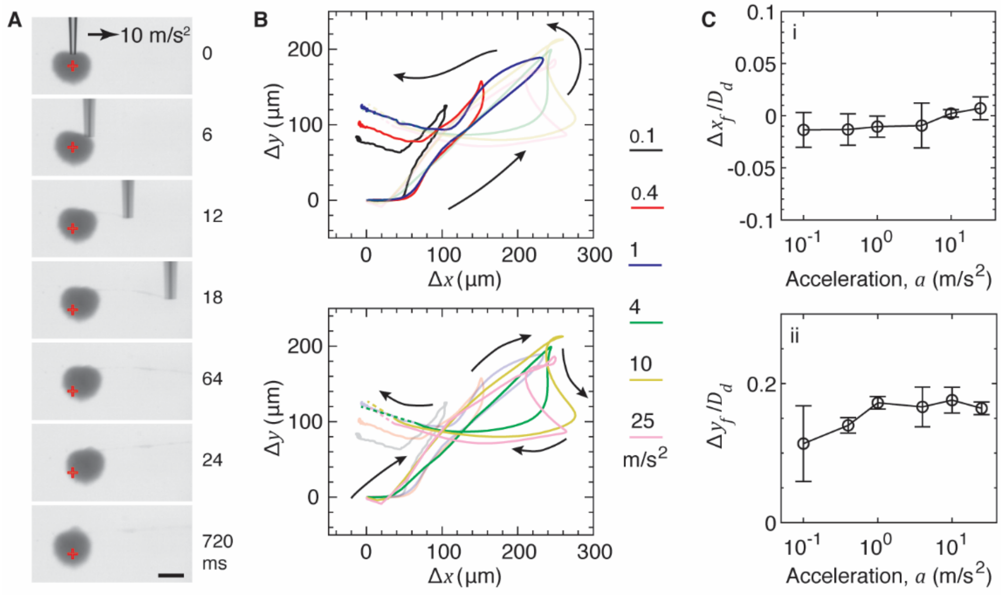


**Figure S5. Using tapered capillary nozzle decreases the final displacement of droplets.** (**A**) A representative time-series of photographs for detaching a tapered capillary nozzle from a droplet at acceleration $a$ =10 m/s$^2$. Nozzle tip diameter, $D_n$ = 120 µm; droplet diameter, $D_d$ = 720 µm. Red cross: center of the droplet at generation stage. Scale bar, 500 µm. (**B**) Trajectories of droplets printed at various accelerations. The nozzle diameter is fixed at $D_n$ = 120 µm, and the droplet diameter is fixed at $D_d$ = 720 µm. Dashed lines: extrapolated trajectories based on the final position of the droplet. (**C**) The final displacement of droplets in (**B**) relative to the droplet diameter $D_d$ at various nozzle acceleration $a$: (i) the displacement along the moving direction of nozzle, $\Delta x_f$; (ii) the displacement along the nozzle axis, $\Delta y_f$. Error bar: STD, *n=3*.
